# Universal assay for measuring vertebrate telomeres by real-time quantitative PCR

**DOI:** 10.1101/797068

**Authors:** Stephanie F. Hudon, Esteban Palencia Hurtado, James D. Beck, Steven J. Burden, Devin P. Bendixsen, Kathleen R. Callery, Jennifer S. Forbey, Lisette P. Waits, Robert A. Miller, Julie A. Heath, Ólafur K. Nielsen, Eric J. Hayden

## Abstract

The study of telomere dynamics in health and aging has been facilitated by qPCR-based telomere length measurements. The widespread application of this approach in vertebrates has been limited by the challenge of developing appropriate reference primers in organisms with genomes that have not been sequenced. Here we report reference primers to highly conserved DNA elements and validate their suitability for qPCR-based telomere measurements across the vertebrate tree of life.

---

Telomeres are the structures at the end of chromosomes that are comprised of proteins bound to repetitive DNA sequences. Telomeres protect the ends of linear chromosomes and provide several important cellular functions^1^. The length of telomere DNA shortens at each cell division, and telomere shortening eventually leads to cellular senescence, which affects tissue function, organismal health and lifespan. Studies in vertebrate systems have shown that the length of telomere repeat regions correlate with phenotypic quality, and can even predict longevity, and reproductive success^2–4^. Telomere lengths are therefore an established predictive biomarker for health and aging. Accelerated rates of telomere shortening have been correlated with multiple causes of stress such as disease, harsh environments, faster growth rates and increased number of offspring^5–8^. Thus, telomere lengths have been used to monitor and predict the effects of different environmental and physiological stressors. The ability to monitor the health and well-bring of organisms and to predict their future success has important applications in captive and wild vertebrates of ecological and economic interest. Understanding telomere dynamics across the vertebrate tree of life could also lead to a better understanding of the mechanisms and evolution of stress responses and aging in general.

Despite this broad potential, the use of telomere length measurements has been limited to relatively few vertebrates, with the majority of studies in avian species. One cause of this limited use is methodological challenges. Real-time quantitative PCR (qPCR) is one of the most widely used methods to measure telomere lengths^9^. Measurement of telomere length by qPCR requires primers that amplify telomere repeats, where the concentration of telomeric DNA in a sample is proportional to telomere length. The primers used to amplify telomeres do not need to be altered for different vertebrate species because the repeating DNA sequence of telomeres is identical in all vertebrates. The assay also requires reference primers that amplify a non-telomeric region of the genome of interest to normalize for the amount of DNA in the sample. However, reference primers optimized in one species may not work in other species, requiring the design and optimization of new primers for each new species. Reference primer development is especially challenging in non-model organisms with genomes that have not been sequenced. As such, the development of primers in species without sequenced genomes is a major bottleneck in the use of telomere length measurements to monitor and predict the health and fitness of vertebrates.

We here report reference primers optimized for qPCR in all vertebrates to broaden the use of telomere length measurements. We designed primer pairs that amplify regions of ultra-conserved elements (UCEs). UCEs are regions of DNA that were originally found to be 100% identical in human, rat and mouse (Fig. 1a)^10^. We predicted that reference primer pairs to these UCEs would be viable in all vertebrates and basal chordates, eliminating the need to design species-specific reference primers for each new telomere assay. Our goal was to find a primer pair for five UCEs that have high amplification efficiency and a single PCR product, as indicated by a single-peak melt curve. We chose UCEs from UCbase 2.0 and designed multiple primer pairs for each using a primer design tool (OligoArchitect Sigma-Aldrich). During design, we limited the product length to 250 bps and designed primers to have a Tm of 60°C to match the Tm of the already developed telomere primers^11^. The top four primer pairs for each of five UCEs, based on software ranking (Beacon Designer™), were synthesized for further testing. We initially tested the primer pairs on mouse (*Mus musculus*) DNA. We used melt-curve analysis to evaluate the specificity of each primer pair and chose a primer pair that gave a single melt peak (Fig. 1b). Amplification efficiencies of each primer pair were determined by performing a serial dilution of target DNA. We found an acceptable primer pair for all five UCEs that showed efficiencies within the best-practice range of 90-110% and correlation values (R^2^) of standard curve dilution replicates that were greater than or equal to 0.98^12^ (Fig. 1c). We chose one primer pair for each UCE for further investigation (Supplementary Table 1).

**Fig. 1.**
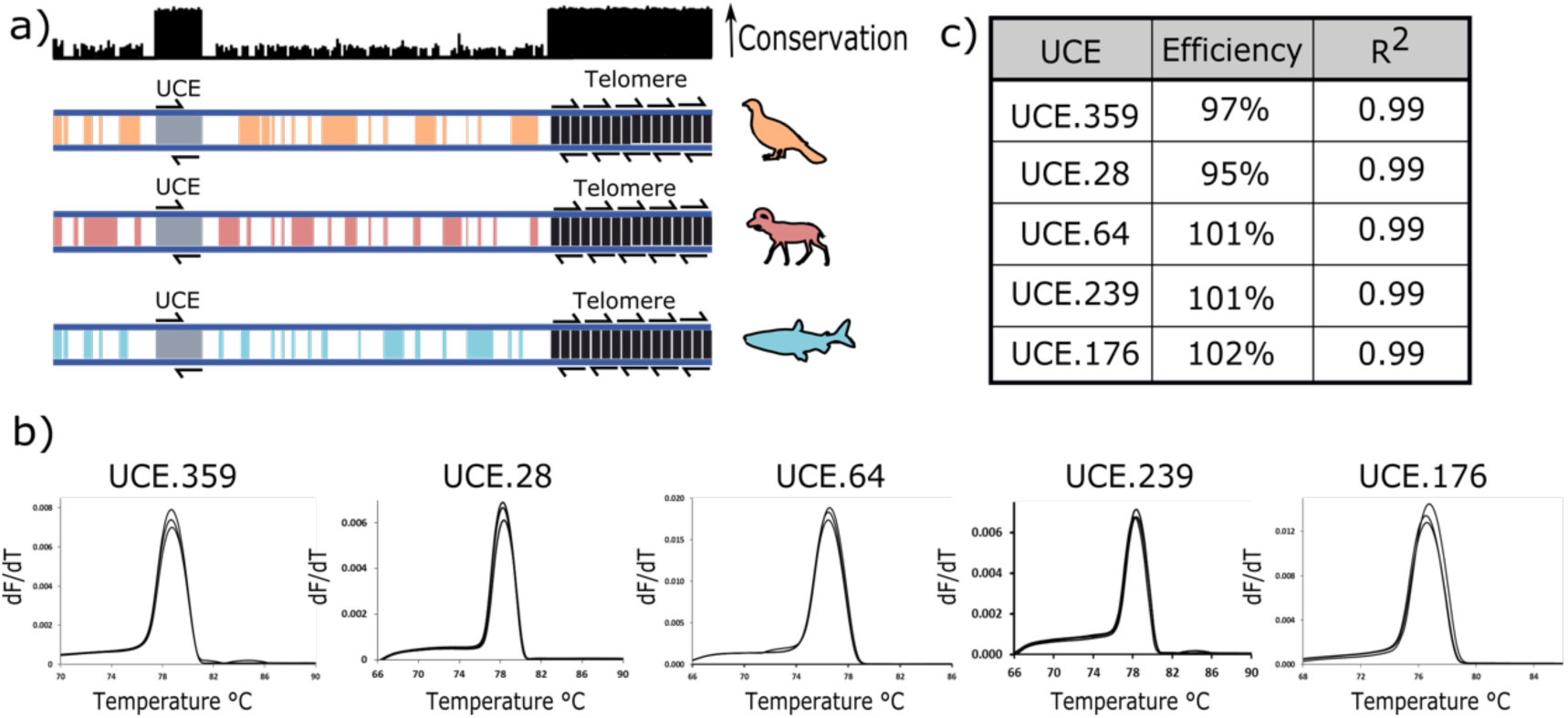
Primer design and quality control. **a**, Primers were designed to ultra-conserved elements which demonstrate conservation across vertebrate genomes. **b**, Melt curves of the chosen UCE primers from qPCR assays in mice. **c**, Efficiencies and R^2^ values of each optimal UCE primer set.

We next validated our primers across the vertebrate tree of life (Fig. 2a). We collected tissue from 19 species, including one basal chordate (sea squirt, *Botryllus schlosseri*) for DNA extraction. We used melt-curve analysis to evaluate the specificity of each primer pair in each organism. We found that all five primer pairs amplified DNA from every species with a single melt peak.

**Fig. 2.**
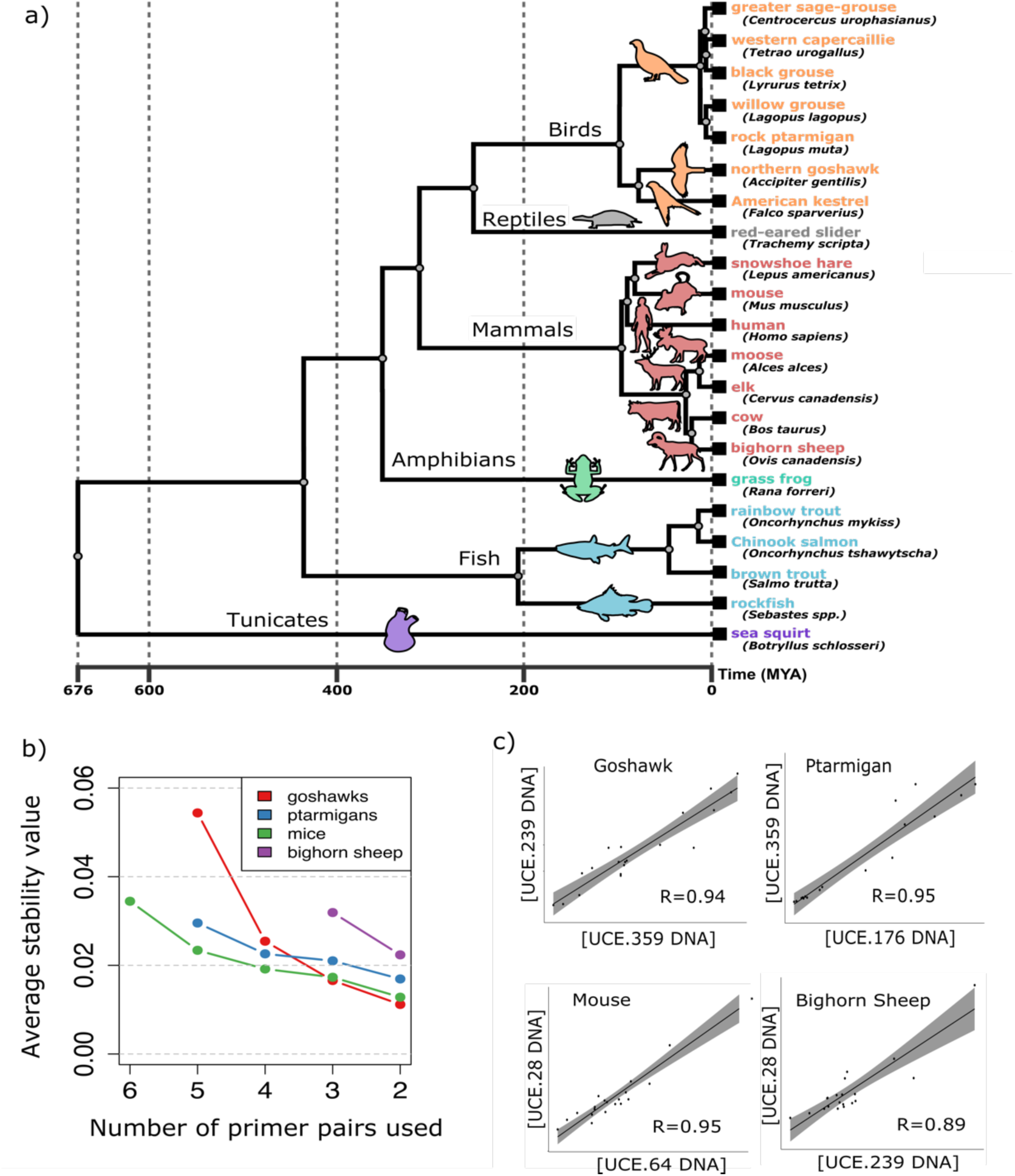
Determining primer pairs across chordates. **a**, Phylogenetic tree of vertebrates (with tunicates as the basal chordate) in which reference primers have been validated. **b**, Average stability values generated in geNorm from the utilization of six to two reference gene primers. **c**, Pearson correlation from the DNA concentrations found using the two best primer pairs for four vertebrate species as determined by geNorm stability values.

An additional requirement for reference genes used for normalizing telomere lengths is that they must not vary in copy number among individuals in the population. For example, a duplication of a reference gene in one individual would appear as a halving of their telomere lengths relative to a non-duplicated individual. The amplification of closely related homolog variants could also introduce normalization differences that would have the same effect as copy number variation. The challenge of choosing the best reference primers for telomere length measurements is similar to choosing primer pairs for normalizing real-time PCR data for gene expression studies, where the goal is to find primers with the least variation among individuals in a population. We therefore evaluated our primer pairs using an algorithm (geNorm) designed for this purpose^13^. The geNorm algorithm evaluates the cumulative variation existing among multiple primer pairs and selects a combination of primer pairs that minimizes the overall variation within the sample array. Two or more primer pairs are preferred over a single primer pair to reduce random experimental variation. The geNorm algorithm iteratively eliminates UCE primer pairs that contribute the most variation until two final primer pairs are chosen. To test this approach, we extracted DNA from 17 to 20 individuals of mouse (*Mus musculus*), rock ptarmigan (*Lagopus muta*), and northern goshawk (*Accipiter gentilis*), representing both sexes and a range of ages and tested these with all five primer pairs.

Using the geNorm package in R, we identified the best combination of UCE primer pairs for each of the three tested species (Supplementary Fig. 1). We found that the use of two primer pairs for normalization leads to very low stability values in each of the species (Fig. 2b,). Pearson correlation of DNA concentrations of the two primers identified through geNorm showed close correlation in these three species (Fig. 2c). Next, we extracted DNA from another mammal (bighorn sheep, *Ovis canadensis*) and tested the top three primer pairs identified in mouse. Using geNorm we again found low stability values for two of the primer pairs. We measured a slightly lower correlation between the best two primers for bighorn sheep than other vertebrate species, which is likely due to the low DNA concentration of these samples (Fig. 2c). However, we anticipate that these three primers would be a good starting place for any ungulate system. For investigators that wish to study telomeres in a new organism, we recommend first testing all five UCE primer pairs on at least ten individuals and performing a stability analysis. Low stability values support using the average DNA concentration for the best two primers to normalize telomere ratios.

We also compared our best two UCE primer pairs for mouse to primer pairs previously reported in the literature for qPCR-based telomere assays in mice^14^. The previously reported primers target the acidic ribosomal phosphoprotein PO (36B4) gene and have been used in multiple publications. We found that our UCE primer pairs are an improvement over the 36B4 primers because they have a single melt-peak while the 36B4 primers have wide multimodal melt peaks which indicates mis-priming or other PCR artifacts (Supplementary Fig. 2). We also note that the 36B4 primer pair was the second primer pair eliminated by the geNorm software, indicating that our top primer pairs are an improvement with respect to stability values in mice. These results suggest that our UCE reference primers may be useful even in organisms with established reference primers.

Telomere length measurements are an important tool for monitoring the health, age and demographics of vertebrate populations. Because telomere lengths have been shown to predict lifespan and reproductive success, telomere lengths may be considered a proxy for fitness. For organisms of conservation concern, telomere lengths have been shown to be an early indicator of extinction risk^15^. The UCE primer pairs reported here can facilitate robust measurements of telomere lengths in vertebrate study systems, even when limited or no genomic sequence data is available. It is important to establish appropriate tissue collection protocols for each new organism because telomere dynamics vary between tissue types^16^. In addition, the biological relevance of shorter telomere lengths in each organism requires knowledge of other health, age and demographic parameters. Nevertheless, understanding telomere length dynamics across taxa increases the number of tools that can be used to address questions about how organisms respond to stressors and will lead to a better general understanding of the evolutionary, environmental and mechanistic aspects of stress and aging across the tree of life.

## Acknowledgement

S.F.H., E.P.H., J.D.B., S.J.B., D.P.B. and E.J.H. were funded by two NSF awards #1826801 and #1807809, as well as an SRC #2018-SB-2842 and NASA award #80NSSC17K0738. J.S.F. was funded by an NSF EPSCoR OIA award #1826801 and K.R.C. and J.A.H. were supported by an NSF REU award DBI: 1263167 and DEB: 1145552. We would also like to thank Boise State University’s Raptor Research Center, the Department of Biological Sciences, College of Arts and Sciences, and Division of Research, American kestrel nest box adopters, and landowners who have nest-boxes on their property as well as the Minidoka Ranger District of Sawtooth National Forest. We acknowledge Jack Connelly (for sage-grouse), Örjan Johansson and Gier Rune Rauset (for capercaillie and black grouse), Rolf Brittas and Tomas Willebrand (for willow grouse), the Icelandic Institute of Natural History and Northeast Iceland Nature Centre (for rock ptarmigan), Knut Keiland (for snowshoe hares), Henrik Andrén and the Grimsö Wildlife Research Station (for moose).

