## Supplementary for "Universal assay for measuring vertebrate telomeres by real-time quantitative PCR" (from another version of this preprint: version 1, 2019-10-09)

### Methods

#### Genomic DNA extraction

DNA was extracted from blood or tissue samples using the Zymo Quick-DNA Microprep Plus Kit (#D4074) according to the manufacture's protocols for the sample type. Blood samples from Northern goshawk and American kestrel were stored in Queen's lysis buffer (0.01 M Tris, 0.01 M NaCl, 0.01 M EDTA, and 1% n-lauroylsarcosine, pH 7.5) prior to extraction. Human DNA was from a buccal swab. DNA from a red-eared slider was extracted from a shell fragment. DNA from sea squirt was extracted from a whole organism. All other DNA samples were extracted from muscle. All animals were treated in accordance with Boise State University animal care and use policies where applicable. DNA purity was assessed by 260/280 nm absorbance ratio, (NanoDrop). DNA was quantified by absorbance at 260 nm and the concentration was normalized to 2ng/µl in an elution buffer containing 10 mM Tris-HCl pH 8.5 and 0.1 mM EDTA (Zymo).

#### Quantitative PCR

Quantitative PCR (qPCR) was performed using a Roche LC96. PCR reactions were carried out in 20 µL volumes containing ~8ng DNA, 10µl of 2x Biotium Fast Plus EvaGreen® qPCR Master Mix and 10 pmol each forward and reverse primers (500 nM final primer concentration). A standard curve for each primer pair was included in triplicate on each plate. The standard curve was prepared by seven serial dilutions (1:4) of DNA. For UCE primers, the two-step thermal cycling profile was 95°C for 2 min, followed by 40 cycles of 95°C for 5 s and 55°C for 30 s, with signal acquisition at the end of the 55°C step, followed by a melt curve generated by increasing temperatures at the end of the thermal cycling. Reference primers were designed to all run with the same thermal cycling profile as the previously characterized telomere primers. DNA concentrations were generated from the standard curve and the ratio of telomere DNA to the average of the two best UCE primers was used to determine relative telomere lengths. All samples were run in triplicate along with three negative controls per primer pair.

### Supplementary

**Table 1**: UCE and Telomere Primer Sequences

| name | Sequence |
|---|---|
| tel 1b | CGGTTTGTTTGGGTTTGGGTTTGGGTTTGGGTTTGGGTT |
| tel 2b | CAGCCGAAAGGCCCTTGGCAGGAGGGCTGCTGGTGGTCTACCCTT |
| UCE.359-F | ATCTGAGACTTGTGACAT |
| UCE.359-R | GTGTTAATTGGTAATGACTATT |
| UCE.28-F | AAATACCACCCAACAGTT |
| UCE.28-R | AAGCCCTATACAGATGGAT |
| UCE.64-F | GAGTCTCCAATATCATCAGAAGC |
| UCE.64-R | ACACATGCCACGATCAATG |
| UCE.239-F | TCAGATGTTCAGCCTATT |
| UCE.239-R | AATACCATGTTAATTATCCTCAA |
| UCE.176-F | TTTCTACAGTTCTGATTTAGTTGA |
| UCE.176-R | TGTTCCCTGTCGCATTAG |

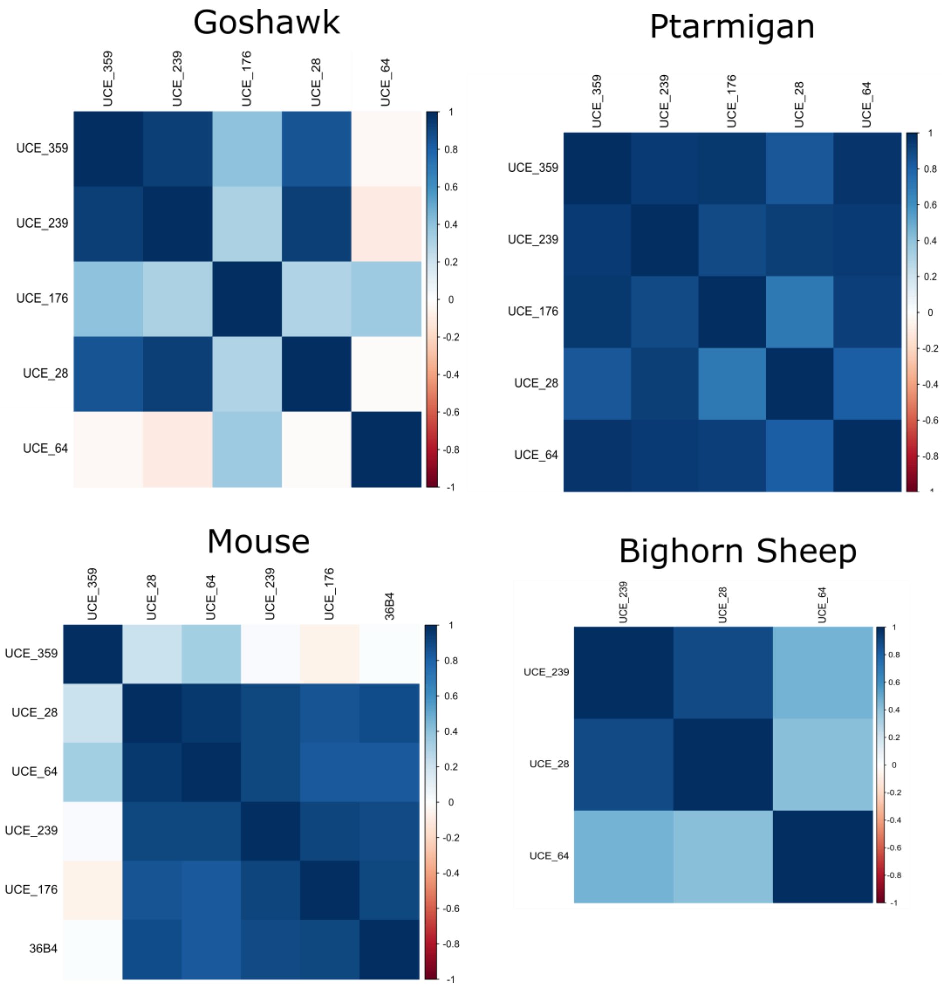


**Supplementary Figure 1**
Pearson Correlation matrixes for UCE primers in four organisms

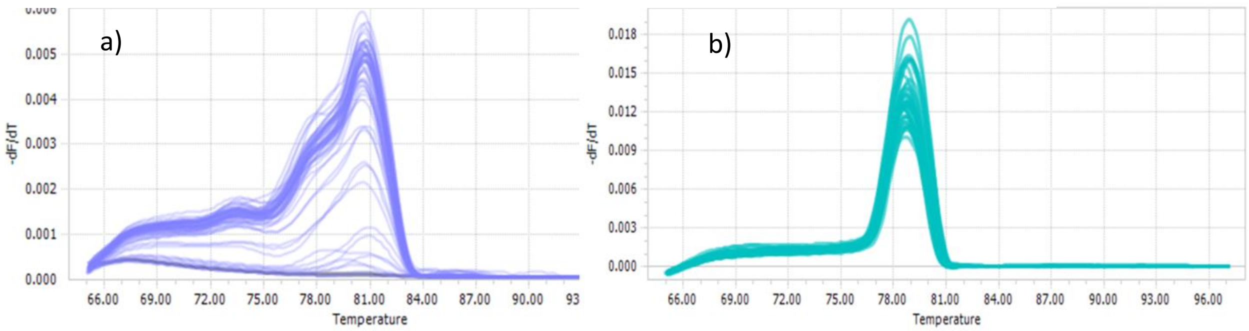


**Supplementary Figure 2**
**a,** Melt curve of 36B4 reference primers used in qPCR-based telomere assays in mice. **b,** Melt curve UCE reference primers in mouse.
